# The adherens junction proteins α-catenin, vinculin and VASP cooperate to promote actin assembly

**DOI:** 10.1101/2022.12.04.518837

**Authors:** Rayan Said, Hong Wang, Julien Pernier, Hemalatha Narassimprakash, Stéphane Roméro, Alexis M. Gautreau, René-Marc Mège, Christophe Le Clainche

## Abstract

The cohesion of tissues requires that cells establish cell-cell junctions. Cells contact each other by forming Arp2/3-dependent lamellipodia before they initiate the formation of cadherin-based adherens junctions (AJs). Maturing AJs then assemble actin under force though the formation of a mechanosensitive complex comprising the actin-binding proteins α-catenin, vinculin and VASP, which individually act on the nucleation, elongation and organisation of actin filaments in different ways. However, the activity of the ternary complex that these actin-regulatory proteins form has not been investigated due to the difficulty of assembling this complex in vitro in the absence of force. Here, we first designed mutants of these proteins that interact independently of force. We then studied their activity by combining actin polymerization kinetics in fluorescence spectroscopy with observation of single actin filaments in TIRF microscopy. Our results reveal how α-catenin, vinculin and VASP combine their activities in a complex to inhibit Arp2/3-mediated branching, stimulate the nucleation and elongation of linear actin filaments from profilin-actin and crosslink these filaments into bundles. These findings shed light on the molecular mechanisms by which actin regulators synergistically control the transition of actin architecture and dynamics that accompanies the formation and maturation of AJs.

## Introduction

The formation and dynamics of tissues during the development of multicellular organisms, their constant repair during wound healing and tissue regeneration, and their adaptation to mechanical perturbations in adults, require adaptative adhesion between cells. Among the cell-cell adhesion structures, cadherin-dependent adherens junctions (AJs) contribute significantly to tissue integrity and maintenance (Collins and Nelson, 2015; Guillot and Lecuit, 2013; Takeichi, 2014). Their action is not limited to passive adhesion since they also act as mechanosensors that trigger adaptation mechanisms to variations in the forces transmitted throughout tissues (Hoffman and Yap, 2015; Ladoux and Mège, 2017; Lecuit and Yap, 2015).

The formation of AJs requires cells to emit membrane protrusions, dependent on the formation of branched actin networks by the Arp2/3 complex, for their cadherins to make contact (Kovacs et al., 2002; Verma et al., 2004; Le Clainche and Carlier, 2008). Once formed, AJs mature into stable structures that assemble an actin network whose organization and dynamics is influenced by the tension that results from the recruitment of myosin II (Heuzé et al., 2019; Mège and Ishiyama, 2017). AJ repair also requires Arp2/3-dependent protrusive activity to bring neighbouring cell membranes together and promote cadherin binding (Li et al., 2021). Deciphering the molecular mechanisms that govern the transition between actin networks of different dynamics and architecture is key to understand the formation and permanent remodelling of AJs. However, the molecular mechanisms underlying actin dynamics associated to AJs is poorly understood.

Cadherins are coupled to the actin cytoskeleton via the interconnected proteins β-catenin, α-catenin, vinculin and VASP. α-catenin has a major structural and regulatory role since its N-terminal dimerization domain dissociates into a monomer that interacts with β-catenin, itself associated with the cytoplasmic domain of cadherins, its central part includes a vinculin binding domain (VBD), and the C-terminal actin-binding domain (ABD) binds to actin filaments (Kobielak and Fuchs, 2004). Along this force transmission pathway, the α-catenin-vinculin interaction is the main mechanosensitive switch that responds to myosin II activity. The actomyosin force transmitted to the ABD domain of α-catenin induces vinculin recruitment to AJs by releasing the autoinhibition of the VBD domain in α-catenin, leading to the reinforcement of AJs (le Duc et al., 2010; Seddiki et al., 2018; Thomas et al., 2013; Yonemura et al., 2010). Single-molecule experiments, using magnetic tweezers and AFM, showed that stretching the central fragment of α-catenin reveals a unique cryptic vinculin binding site (VBS) which reversibly binds vinculin at 5 pN (Yao et al., 2014; Maki et al., 2018). Although the binding of β-catenin to α-catenin breaks the α-catenin dimer and reduces its affinity for actin filaments in solution (Drees et al., 2005), subsequent manipulations of single molecules with optical tweezers and structural studies showed that β-catenin-α-catenin heterodimers bind actin under force through a catch bond with an optimum at a force of 5 pN (Buckley et al., 2014; Mei et al., 2020; Xu et al., 2020). Finally, the actin regulator VASP interacts with a FPPPP motif in the proline-rich linker of vinculin (Brindle et al., 1996).

Several lines of evidence suggest that the proteins that link cadherins to actin, mentioned above, also regulate actin polymerization in a force-dependent manner. Indeed, the knock-downs of non-muscle myosin II isoforms (NMIIA and NMIIB) cause a decrease in the amount of junctional F-actin and barbed end density at AJs, suggesting the existence of an actin nucleation mechanism dependent on actomyosin tension (Leerberg et al., 2014). This activity involves the recruitment of vinculin to AJs through its force-dependent interaction with the vinculin binding site (VBS) of α-catenin. This mechanism also requires the binding of Mena/VASP to the FPPPP motif of the proline-rich linker of vinculin (Leerberg et al., 2014; Vasioukhin et al., 2000). These data suggest that the force-dependent formation of a ternary complex containing α-catenin, vinculin and VASP induces actin assembly though an unknown mechanism. At the molecular level, α-catenin, vinculin and VASP possess ABDs that act on actin polymerization and organisation in different ways. Vinculin, via its actin filament binding domain Vt, nucleates actin filaments capped at their barbed ends and bundles these filaments (Le Clainche et al., 2010). The bundling activity of vinculin requires the dimerization of Vt (Thompson et al., 2017). α-catenin possesses barbed end capping activity through its ABD domain (Hansen et al., 2013). Finally, tetrameric proteins of the VASP family cumulate nucleation, barbed end elongation, and bundling activities (Breitsprecher et al., 2011, 2008; Hansen and Mullins, 2010; Laurent et al., 1999).

The individual activities of the AJ proteins on actin polymerization detailed above do not allow to predict the relative importance of these proteins, nor the activity that results from their combination in AJ dynamics. Here, we used mutated versions of the proteins that interact constitutively without the action of a mechanical stimulus to reconstitute the α-catenin-vinculin-VASP complex and determined its activity by combining kinetic measurements in fluorescence spectroscopy with quantification of the polymerization and organisation of actin filaments by microscopy. Our results reveal the mechanism by which α-catenin, vinculin and VASP combine their activities to inhibit Arp2/3-mediated branching, enhance actin filament nucleation and barbed end growth from profilin-actin, and bundle these filaments.

## Results

### α-catenin, vinculin and VASP combine their activities to stimulate actin assembly in the presence of profilin

To determine whether and how the machinery composed of the proteins α-catenin, vinculin and VASP regulates actin polymerization, we used purified recombinant proteins (Supplementary Figure 1). The formation of a complex containing these three proteins is not constitutive because, in α-catenin, the VBS is masked by the interaction of the M1 α-helix bundle that contains it with the adjacent α-helix bundles MII and MMIII. Similarly, vinculin is auto-inhibited by a well-documented intramolecular interaction between vinculin head (Vh), which interacts with the VBS of several proteins including α-catenin, and vinculin tail (Vt), which interacts with actin filaments. We therefore used an α-catenin construct called Δmod, deleted from the MII and MIII domains, that constitutively exposes the VBS domain (Seddiki et al., 2018). We also used a vinculin construct called V_1ab4_ containing point mutations in the Vh domain that weakens its autoinhibition, allowing its interaction with the VBS of α-catenin (Wang et al., 2022)(Supplementary Figure 1). As a first step, we determined the actin polymerization activity of these proteins by measuring the polymerization kinetics of pyrene-labelled actin in fluorescence spectroscopy (Figure 1). This method allows the detection of small variations and the rapid screening of multiple conditions before the direct observation of actin filaments in fluorescence microscopy. Because physicochemical conditions in the vicinity of a membrane vary due to ionic fluxes, enrichment with charged lipids such as PIP2, or molecular crowding with proteins of different charge, we varied the ionic strength conditions of the assays from 25 mM KCl to 100 mM KCl. Of the three proteins tested alone, only VASP had significant actin polymerization stimulating activity, while α-catenin (WT and Δmod) and Vinculin V_1ab4_ showed little or no activity at 25 mM KCl (Figure 1A, B). The pairwise association of the proteins showed that V_1ab4_, which is inactive alone, increased the activity of VASP at 25 mM KCl in a dose dependent manner (Figure 1A, B, C, Supplementary Figure 2A). Similarly, the combination of V_1ab4_ and Δmod resulted in stimulation of actin assembly, which appeared to be the result of the exposure of the Vt domain of V_1ab4_ following its activation by Δmod (Figure 1B, D, Supplementary Figure 2B). This hypothesis is supported by the fact that the activity was not observed with WT α-catenin, which does not bind vinculin (Figure 1A). The fact that VASP and α-catenin/Δmod, which are not known to interact directly, stimulated actin assembly in a synergistic manner was not expected, but suggests that these two ABPs cooperate when combined in the same protein complex. Finally, we observed the strongest stimulation of actin assembly in the presence of all three proteins (Figure 1A, B). Increasing ionic strength reduced the ability of the three proteins to stimulate actin assembly. The maximum effect was observed at 25 mM KCl, while the activity remained strong at 50 mM KCl, which is the ionic strength used in many studies, and little effect was observed at 100 mM KCl (Figure 1E, Supplementary Figure 2C).

**Figure 1.**
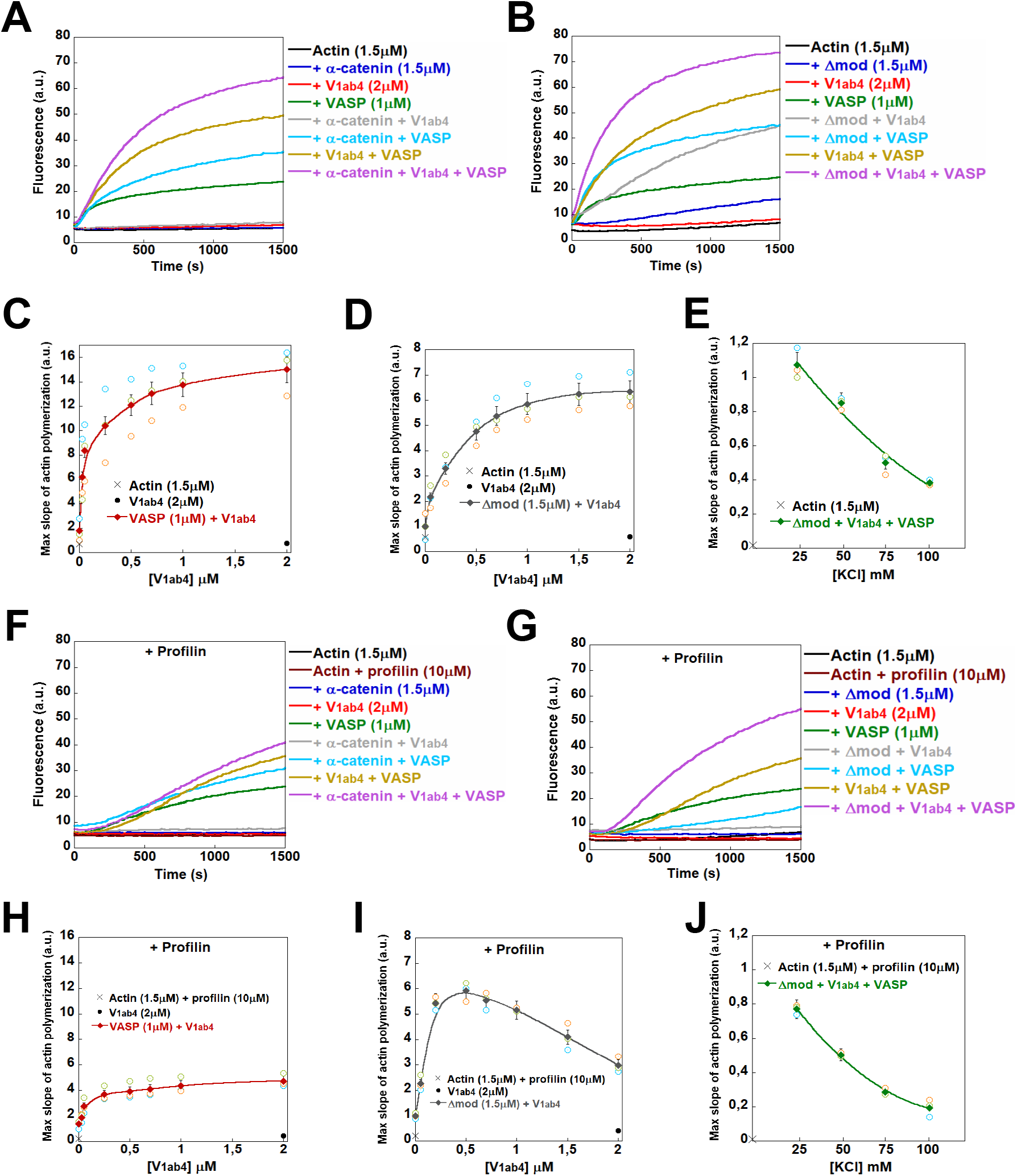
α-catenin, vinculin and VASP cooperate to stimulate actin polymerization in the presence of profilin. **(A,B)** Spontaneous nucleation of 1.5 μM G-actin (10% pyrene-labeled) was measured in the presence of the indicated combinations of 1.5 μM α-catenin, 2 μM V_1ab4_, 1 μM VASP **(A)**, or 1.5 μM Δmod, 2 μM V_1ab4_, 1 μM VASP **(B)** in a low salt buffer (25 mM KCl). **(C, D)** The maximal rate of spontaneous actin polymerization (1.5 μM G-actin, 10% pyrene-labeled) is plotted as a function of increasing concentrations of V_1ab4_ in the presence of 1 μM VASP **(C)** or 1.5 μM Δmod **(D)**. **(E)** The maximal rate of spontaneous actin polymerization (1.5 μM G-actin, 10% pyrene-labeled) is plotted against increasing concentrations of KCl in the presence of 1.5 μM Δmod, 2 μM V_1ab4_, 1 μM VASP. **(F,G)** Spontaneous nucleation of 1.5 μM G-actin (10% pyrene-labeled) was measured in the presence of 10 μM profilin and the indicated combinations of 1.5 μM α-catenin, 2 μM V_1ab4_, 1 μM VASP **(F)**, or 1.5 μM Δmod, 2 μM V_1ab4_, 1 μM VASP **(G)**, in a low salt buffer (25 mM KCl). **(H, I)** The maximal rate of spontaneous actin polymerization (1.5 μM G-actin, 10% pyrene-labeled) is plotted as a function of increasing concentrations of vinculin V_1ab4_ in the presence of 10 μM profilin and 1 μM VASP **(H)** or 10 μM profilin and 1.5 μM Δmod **(I)**. **(J)** The maximal rate of spontaneous actin polymerization (1.5 μM G-actin, 10% pyrenyl-labeled) is plotted against increasing concentrations of KCl in the presence of 1.5 μM Δmod, 2 μM V_1ab4_, 1 μM VASP and 10 μM profilin. These experiments were performed three times with the same conclusions. Open symbols with different colors represent three independent experiments in the same conditions, and closed symbols indicate the average of the triplicates. Error bars represent ±SEM (n=3).

Profilin is an essential protein in the mechanisms governing actin assembly in cells, as it is generally accepted that the majority of the polymerizable cytoplasmic actin pool is complexed to profilin. One of the main functions of profilin is to prevent spontaneous nucleation of actin in the cytoplasm so that it can be controlled in time and space by specific machineries. In addition, profilin serves as a co-factor for many actin regulatory proteins, including VASP or formins (Breitsprecher et al., 2008; Romero et al., 2004). In agreement, we observed here, in the presence of profilin, the strongest stimulation of actin assembly in the presence of all three proteins at 25 mM KCl (Figure 1F, G).

These experiments also confirmed that α-catenin Δmod, with an exposed VBS, conferred more activity than α-catenin WT (Figure 1F, G). Profilin did not abolish the ability of V_1ab4_ to synergize with VASP and Δmod (Figure 1H, I, Supplementary Figure 2D, E). Our data suggest that VASP is likely the protein that plays the most important role in inducing actin polymerization in the presence of profilin, as it alone possesses this activity, whereas the role of the other proteins seems to be to increase the activity of VASP (Figure 1F, G). Increasing the ionic strength in the presence of profilin also reduced the ability of all three proteins to stimulate actin polymerization. The maximum effect was observed at 25 mM KCl, remained strong at 50 mM KCl, and was greatly reduced at 100 mM KCl (Figure 1J, Supplementary Figure 2F).

As the kinetics of spontaneous actin assembly can reflect a concomitant stimulation of nucleation and elongation, we decided to uncouple these two activities by adding spectrin-actin seeds in our reactions, which boosts elongation and makes the spontaneous nucleation of actin a negligible parameter (Figure 2). To avoid the contribution of V_1ab4_ and VASP nucleation activities, which are favoured at low ionic strength, the experiments were performed only at 100 mM KCl. In these conditions, VASP effectively protected against the capping activity of V_1ab4_ and α-catenin (Figure 2A, C, Supplementary Figure 3A), but only modestly protected against the Δmod capping activity (Figure 2B), suggesting that the absence of the MII-MIII domains in Δmod leads to a better exposure of the C-terminal ABD of α-catenin that caps actin filament barbed ends. Surprisingly, when the three proteins were added together, they synergised to accelerate elongation at a rate that was greater than that of VASP alone. This observation suggests that VASP protects the barbed ends of actin filaments against the combined capping activity of V_1ab4_ and α-catenin/Δmod, but also that V_1ab4_ and α-catenin/Δmod assist VASP to increase the elongation of these actin filaments (Figure 2A, B). The surprising loss of influence of Δmod capping activity in the presence of V_1ab4_ and VASP could result from a hierarchy between the activities, leading to V_1ab4_ imposing its activity on Δmod. This would explain why Δmod capping activity did not add to that of V_1ab4_ when the 2 proteins were together in the absence of VASP (Figure 2B). Importantly, in the presence of profilin, VASP, combined with V_1ab4_ and α-catenin/Δmod, also promoted barbed-end elongation (Figure 2D-F, Supplementary Figure 3B).

**Figure 2.**
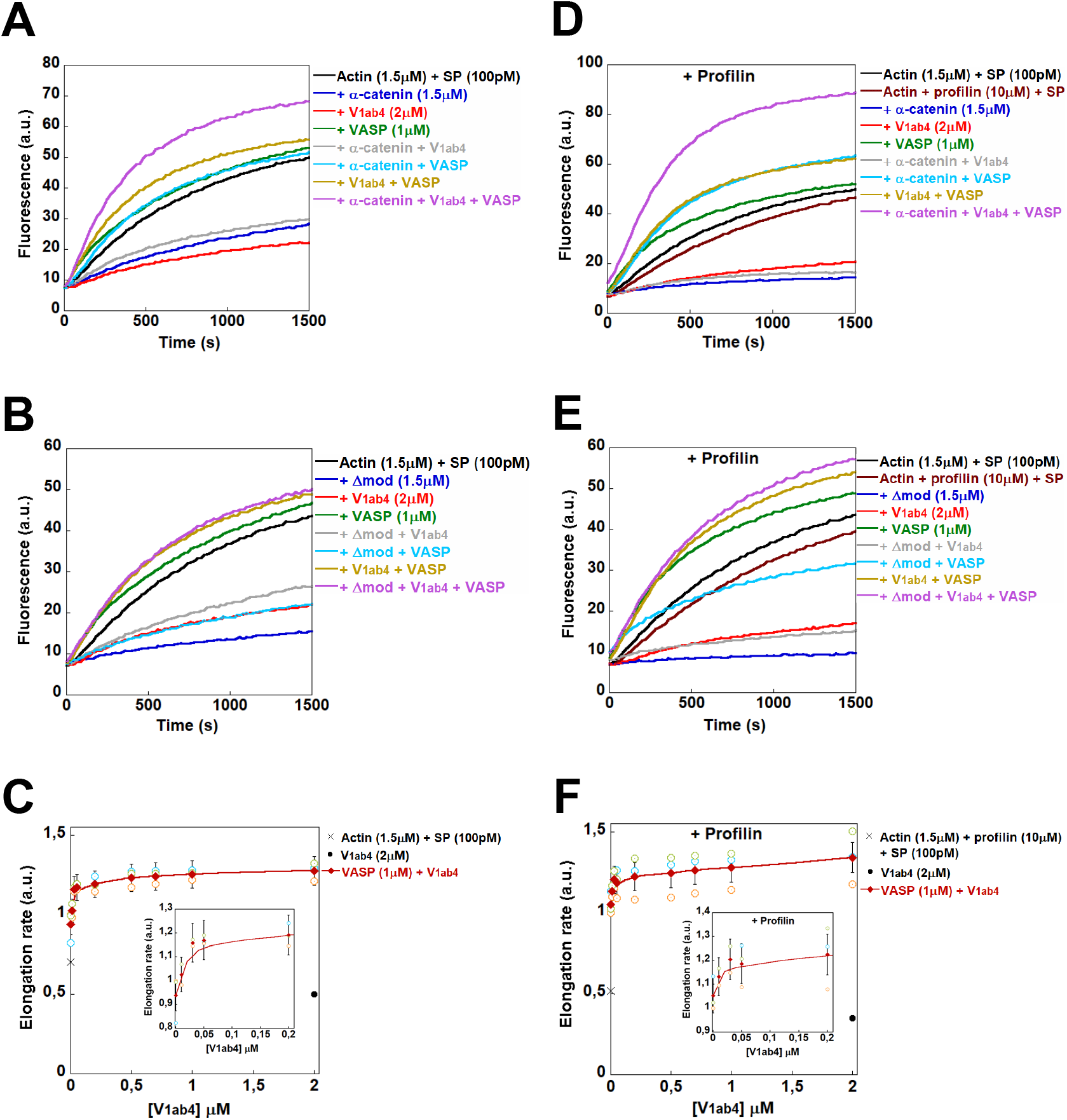
α-catenin, vinculin and VASP synergize to promote barbed-end elongation in the presence of profilin. **(A,B)** Barbed end elongation was measured in the presence of 100 pM spectrin-actin seeds (SP), 1.5 μM G-actin (10% pyrene-labeled) and the indicated combinations of 1.5 μM α-catenin, 2 μM V_1ab4_, 1 μM VASP **(A)** or 1.5 μM Δmod, 2 μM V_1ab4_, 1 μM VASP **(B)** in a high salt buffer (100 mM KCl). **(C)** Barbed end elongation was measured in the presence of 100 pM spectrin-actin seeds, 1.5 μM G-actin (10% pyrene-labeled), 1 μM VASP, and increasing concentrations of V_1ab4_ in a high salt buffer (100 mM KCl). The fraction of barbed end elongation was then calculated as the ratio between the elongation rate in the presence of the indicated proteins and the elongation rate of actin alone. **(D,E)** Barbed end elongation was measured in the presence of 100 pM spectrin-actin seeds, 1.5 μM G-actin (10% pyrene-labeled) in the presence of 10 μM profilin and the indicated combinations of 1.5 μM α-catenin, 2 μM V_1ab4_, 1 μM VASP **(D)**, or 1.5 μM Δmod, 2 μM V_1ab4_, 1 μM VASP **(E)**, in a high salt buffer (100 mM KCl). **(F)** Barbed end elongation was measured in the presence of 100 pM spectrin-actin seeds, 1.5 μM G-actin (10% pyrenyl-labeled), 1 μM VASP and 10 μM profilin, and increasing concentrations of V_1ab4_. The fraction of barbed end elongation was then calculated as the ratio between the elongation rate in the presence of the indicated proteins and the elongation rate of actin alone. These experiments were performed three times with the same conclusions. Open symbols with different colors represent three independent experiments in the same conditions, and closed symbols indicate the average of the triplicates. Error bars represent ±SEM (n=3).

### The stimulation of actin assembly by the combined action of α-catenin, vinculin and VASP results from an increase in both nucleation and barbed end elongation

As the interpretation of kinetic studies of actin polymerization inevitably reaches its limits in the presence of three proteins that combine their individual activities, we observed their effect on single actin filaments in TIRF microscopy to confirm nucleation and barbed-end elongation activities. In this series of experiments, we used the experimental conditions tested previously to determine the density of actin filaments, reflecting actin nucleation, and their elongation rate in the presence of V_1ab4_, Δmod and VASP, alone or in combination, at low (25 mM KCl) and high (100 mM KCl) ionic strength, with and without profilin. We first compared the density of filaments that form in the presence of these proteins over time. Although very few filaments were observed in the presence of the proteins alone, we observed an increase in the density of filaments when the three proteins were combined at both 25 mM and 100 mM KCl (Figure 3A, C, D, Supplementary movie 1, Supplementary movie 2), indicating an effect on nucleation. In the presence of the three proteins, the filaments were also longer, suggesting an effect on elongation (Figure 3A). We also observed, when the three proteins were combined, an increase in the density and length of actin filaments produced in the presence of profilin (Figure 3B, C, D, Supplementary movie 3, Supplementary movie 4).

**Figure 3.**
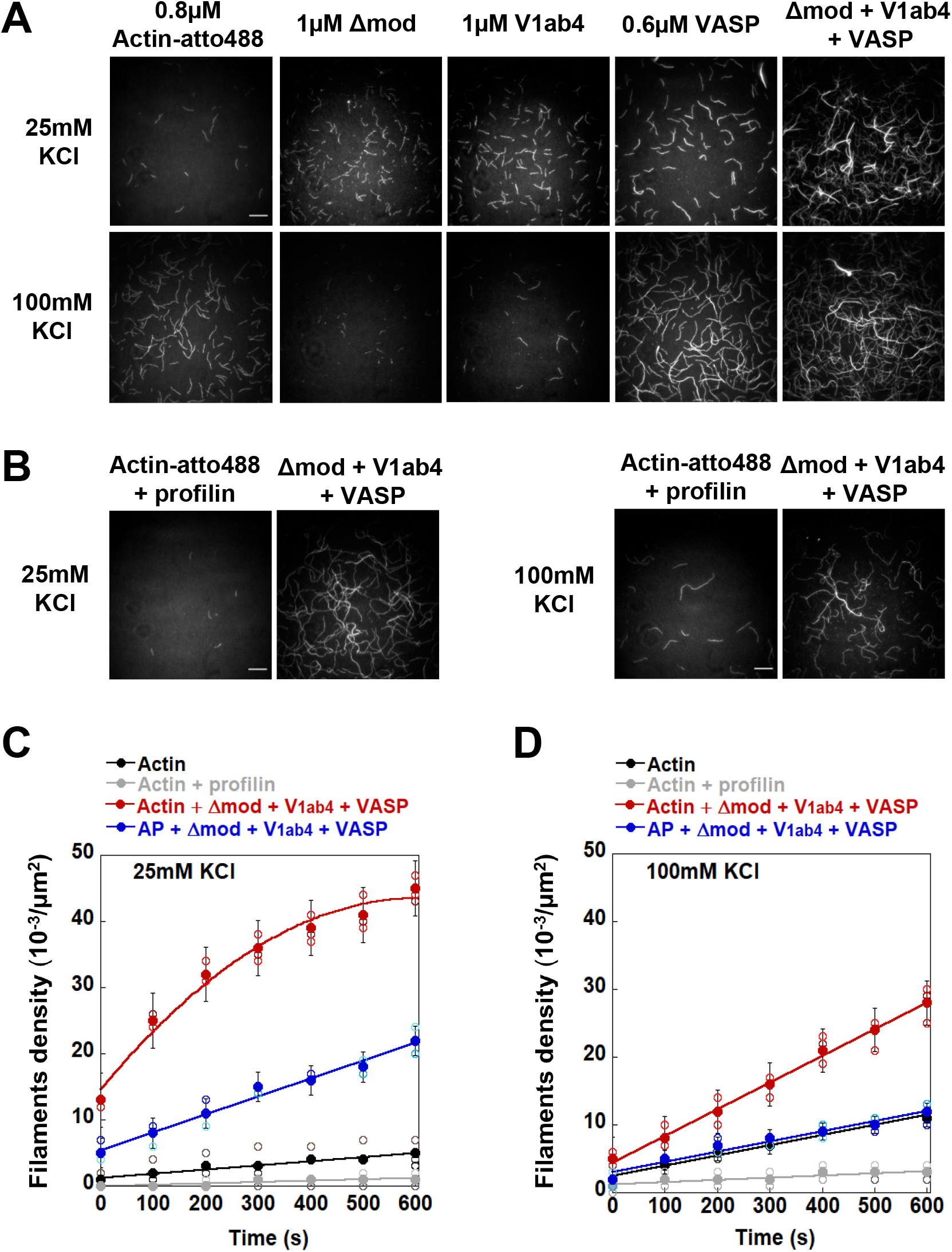
Direct observation of single actin filaments in TIRF microscopy in the presence of Δmod, V_1ab4_ and VASP. **(A)** Single actin filaments observed in TIRF microscopy in the presence of 0.8 μM actin (10% Atto488-labelled) alone and supplemented with the indicated combinations of 1 μM Δmod, 1 μM V_1ab4_ and 0.6 μM VASP, in low salt (25 mM KCl) or high salt (100 mM KCl). See supplementary movies 1 and 2. **(B)** Single actin filaments observed in TIRF microscopy in the presence of 0.8 μM actin (10% Atto488-labeled) and 5 μM profilin, and supplemented with the indicated combinations of 1 μM Δmod, 1 μM V_1ab4_ and 0.6 μM VASP, in low salt (25 mM KCl) or high salt (100 mM KCl). See supplementary movies 3 and 4. Each image represents a complete field. **(A,B)** Scale bar = 15 μm. **(C,D)** Quantification of filament density as a function of time from the experiments shown in (A,B) at 25 mM KCl **(C)** and 100 mM KCl **(D)**. In (C) and (D) AP stands for actin + profilin. These experiments were performed three times with the same conditions. Open symbols with different colors represent three independent experiments in the same conditions, and closed symbols indicate the average of the triplicates. Error bars represent ±SEM (n=3). Images were acquired in 590 s.

From the same experiments, we also measured the elongation rate of the barbed ends of actin filaments (Figure 4). At low ionic strength (25 mM KCl), proteins alone moderately affected filament elongation positively or negatively (Figure 4C). However, their combination resulted in a 2-3 fold increase in the rate of actin filament barbed end elongation, which was a specific activity of the V_1ab4_-Δmod-VASP combination, as it did not correspond to the sum of the activities measured for the isolated proteins (Figure 4A, C). The addition of profilin, under the same low ionic strength conditions, allowed the combination of V_1ab4_, Δmod and VASP to accelerate actin filament elongation by up to a factor of 5 compared to the control with actin alone (Figure 4B, C). Observation of the details of single filament elongation revealed that it was regularly interrupted by marked pauses, reflecting capping events of the barbed ends by V_1ab4_ and Δmod (Figure 4B). These events were rarely observed in the presence of the same combination of proteins without profilin (Figure 4A). At high ionic strength (100 mM KCl), the combination of V_1ab4_, Δmod and VASP also led to the acceleration of actin filament elongation in the absence and presence of profilin (Figure 4D, E). The detailed quantification of these experiments revealed that this elongation activity emerges from the combined action of V_1ab4_, Δmod and VASP and was not the simple sum of the inhibitions and activations observed for the proteins alone (Figure 4F).

**Figure 4.**
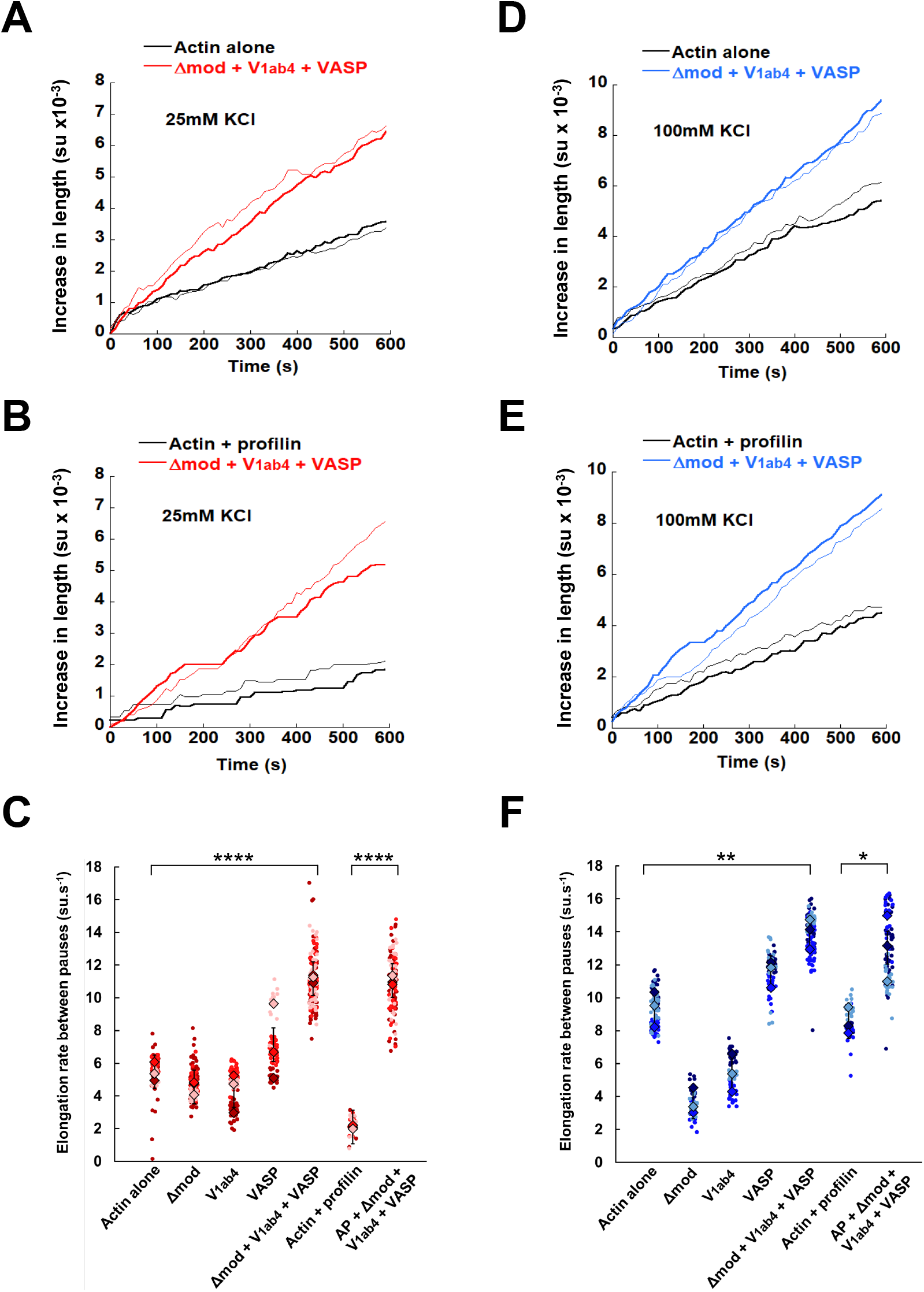
Stimulation of actin filaments elongation by α-catenin, vinculin and VASP. **(A,B)** Traces corresponding to the elongation of two filaments under the indicated conditions (from Figure 3A) in low salt conditions (25 mM KCl) in the absence **(A)** or presence of profilin **(B)**. **(C)** Quantification of the elongation rate of the barbed end of filaments between pauses under the indicated conditions at 25 mM KCl. **(D,E)** Traces corresponding to the elongation of two filaments under the indicated conditions (from Figure 3B) in high salt conditions (100mM KCl) in the absence **(D)** or presence of profilin **(E)**. **(F)** Quantification of the elongation rate of the barbed end of filaments between pauses under the indicated conditions at 100 mM KCl. **(A,B,D,E)** The two traces with same color represent two filaments from two independent experiments in the same conditions. **(C,F)** Closed circle symbols with three different colors represent filaments elongation rate from three independent experiments in the same conditions. The mean of each independent experiment is represented by diamond symbols of three different colors. Error bars represent ±SEM (n=3). A significant difference was found using a two-tailed *t*-test comparing the mean values. ****, *p*<0.0001; **, *p*<0.01; *, *p*<0.05.

Taken together, our direct observations in microscopy confirm the kinetic assays by showing that the three proteins synergize to nucleate actin filaments and accelerate the elongation of their barbed ends in the presence of profilin.

### α-catenin, vinculin and VASP synergise to form actin filament bundles

The TIRF microscopy observations presented above revealed the formation of actin filament bundles when all three proteins V_1ab4_, Δmod, and VASP were present, while no bundles were observed when actin filaments were grown alone (Figure 3A, B). To quantify this activity precisely, we combined kinetics of actin filament bundling measured by light scattering with fluorescence microscopy observations of the same reactions (Figure 5). Light scattering kinetics showed that neither V_1ab4_ nor Δmod alone induced bundle formation, whereas VASP showed low activity (Figure 5A-D). These experiments also revealed synergies between the protein pairs and the three proteins to form actin bundles (Figure 5A-D). Replacement of α-catenin with Δmod, in which VBS is exposed, allowed for a higher bundling activity when combined with V_1ab4_ and VASP (Figure 5C, D), indicating that the α-catenin-vinculin complex contributes to actin filament crosslinking. Direct observation of the reactions in fluorescence microscopy confirmed the formation of sparse bundles by the proteins alone and larger bundles by the protein pairs and the combination of the three proteins (Figure 5E).

**Figure 5.**
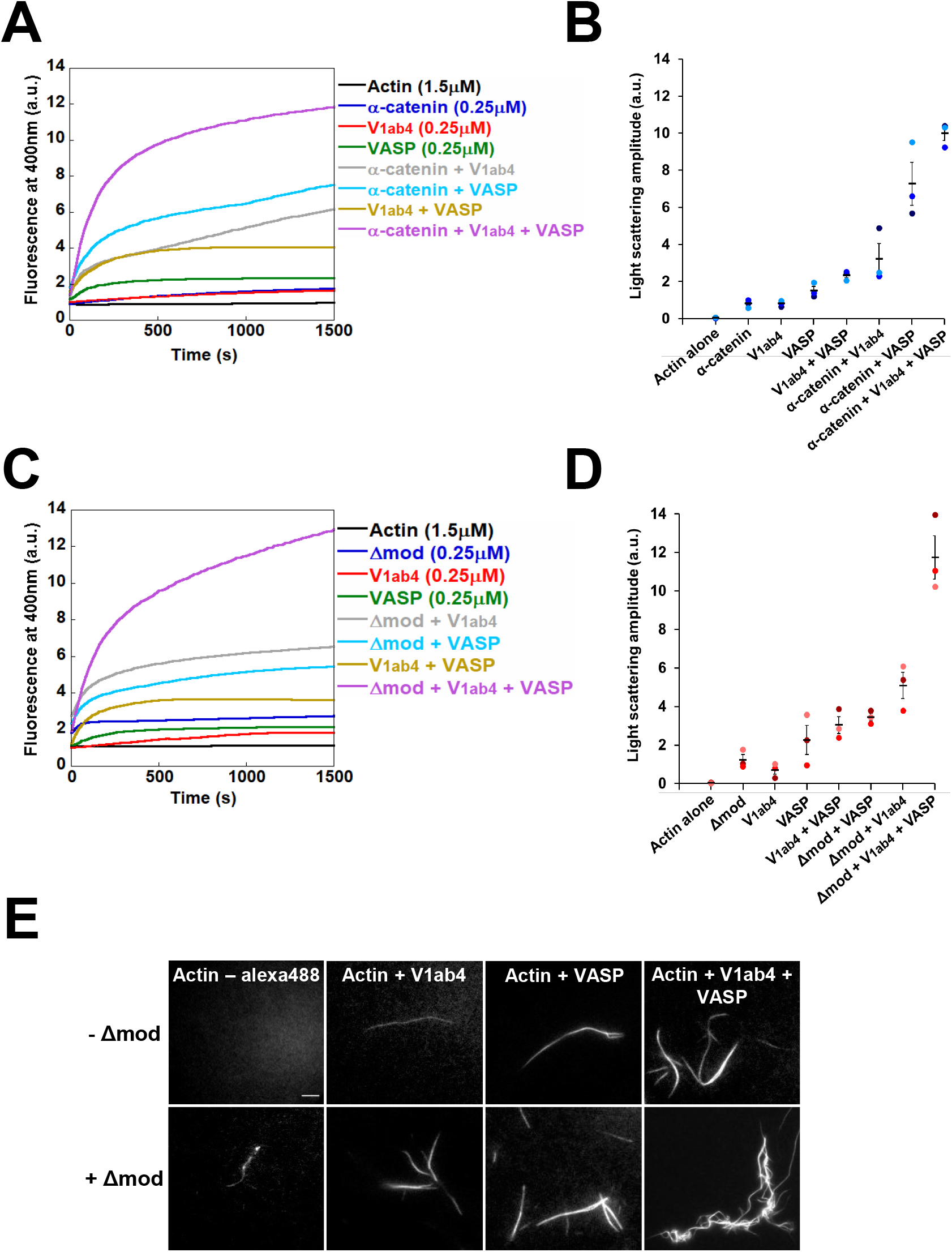
α-catenin, vinculin and VASP bundle actin filaments synergistically. **(A)** Light scattering was measured during the polymerization of actin alone (1.5 μM G-actin), and in the presence of all the combinations of 0.25 μM α-catenin WT, 0.25 μM V_1ab4_ and 0.25 μM VASP, in low salt (25 mM KCl). **(B)** Quantification of light scattering amplitude at 1500 s for each conditions indicated in (A). **(C)** Light scattering was measured during the polymerization of actin alone (1.5 μM G-actin), and in the presence of all the combinations of 0.25 μM α-catenin Δmod, 0.25 μM V_1ab4_ and 0.25 μM VASP, in low salt (25 mM KCl). **(D)** Quantification of light scattering amplitude at 1500 s for each conditions indicated in (C). **(A-D)** These experiments were performed three times with the same conditions. **(B, D)** Symbols with different colors represent three independent experiments. Error bars represent ±SEM (n=3). **(E)** Representative epifluorescence images of Alexa-488 actin filament bundles observed alone and in the presence of all the combinations of Δmod, V_1ab4_ and VASP. Conditions: 1.5 μM G-actin (2%Alexa-488-labeled), 0.25 μM Δmod, 0.25 μM V_1ab4_ and 0.25 μM VASP. Scale bar = 15 μm.

### α-catenin, vinculin and VASP inhibit Arp2/3-mediated branching of actin filaments to promote their cross-linking into fast-growing bundles

Early actin polymerization kinetics experiments showed that α-catenin inhibits the formation of branched actin networks by the Arp2/3 complex (Drees et al., 2005), while more recent studies showed that talin-activated vinculin reorganizes branched actin networks into bundles (Boujemaa-Paterski et al., 2020). These observations led us to test the effect of combinations of α-catenin, vinculin and VASP on the formation and organization of Arp2/3 dependent actin networks. Actin polymerization kinetics (Figure 6A), quantified by measuring the half-time of polymerization (Figure 6B), showed that Δmod and vinculin V_1ab4_ alone inhibited the actin polymerization normally stimulated by the Arp2/3 complex in the presence of the VCA domain of N-WASP. In contrast, VASP alone had little effect on this reaction. The combination of Δmod, V_1ab4_ and VASP stimulated actin polymerization in the presence of Arp2/3 and VCA at a level identical to that stimulated by these three proteins in the absence of Arp2/3 and VCA, suggesting that ΔMod, V_1ab4_ and VASP imposed their combined activity to the branched reaction (Figure 6A, B). The direct observation of this reaction in TIRF microscopy revealed a progressive reorganization of the branched actin network by Δmod, V_1ab4_ and VASP (Figure 6C, Supplementary movie 5). From a densely branched network of filaments in the presence of Arp2/3-VCA, the network became sparse and weakly branched in the presence of Δmod, weakly branched and denser in the presence of Δmod and V_1ab4_, and densely populated with long, unbranched and strongly bundled filaments in the presence of Δmod, V_1ab4_ and VASP (Figure 6C, Supplementary movie 5). Kymographs showed not only the transition from a branched to a bundled network, but also the acceleration of the growth of filaments that develop from the bundles (Figure 6D), recapitulating the actions of Δmod, V_1ab4_ and VASP on the bundling and elongation of filaments.

**Figure 6.**
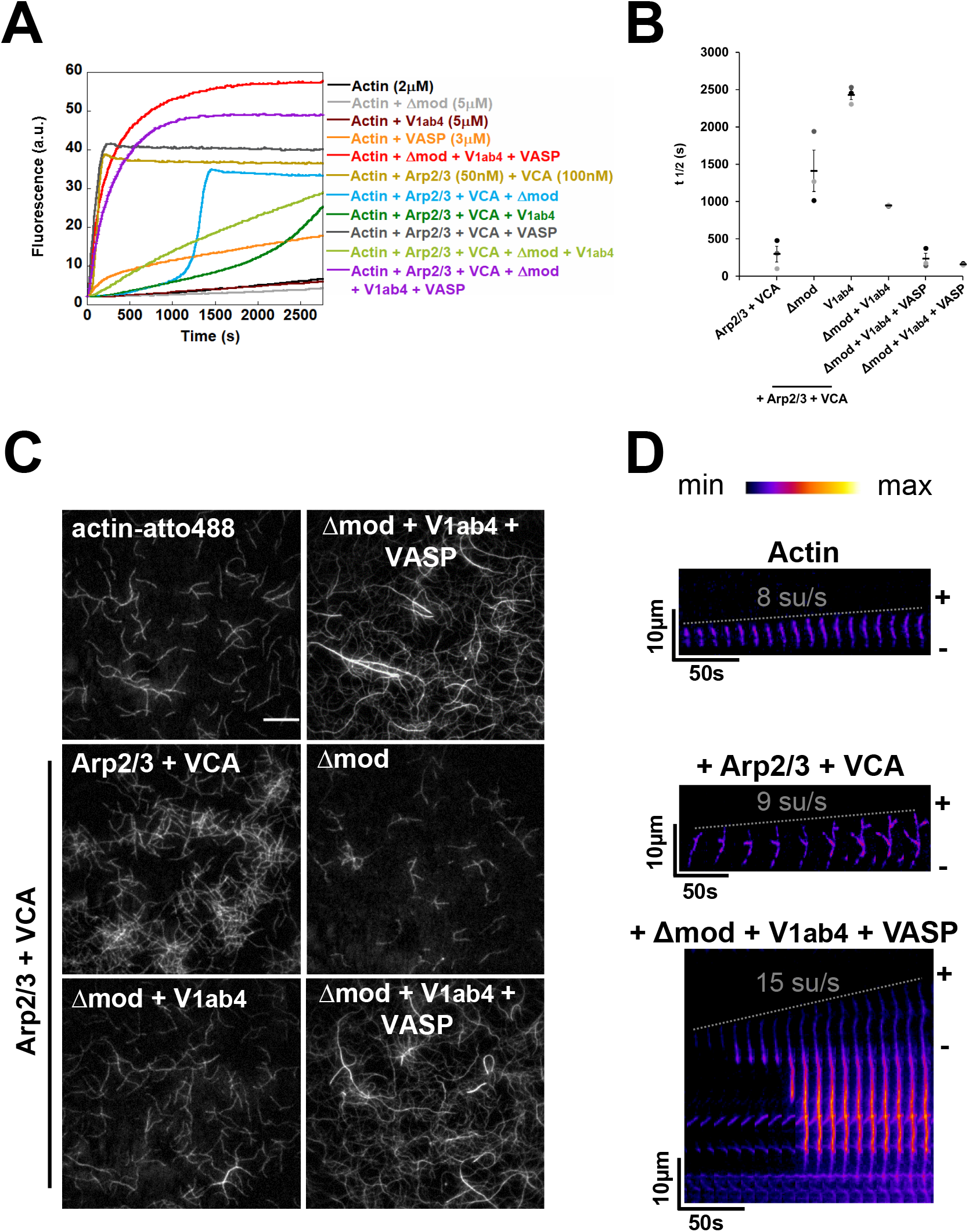
Remodeling of Arp2/3-mediated branched actin into fast-growing bundles by α-catenin, vinculin and VASP. **(A)** Kinetics of actin polymerization were measured in the presence of 2 μM G-actin (10% pyrene-labelled), 50 nM Arp2/3, 100 nM VCA, 5 μM Δmod, 5 μM V_1ab4_ and 3 μM VASP, in conditions that allow nucleation and elongation (50 mM KCl). **(B)** Quantification of the polymerization half-time (t½) for conditions indicated in (A). These experiments were performed three times with the same conclusions. Symbols with different colors represent three independent experiments. Error bars represent ±SEM (n=3). **(C)** TIRF microscopy observation of the transition from actin filaments branching to bundling in the presence of 0.8 μM actin (10% Atto488-labelled), supplemented with the indicated combinations of 50 nM Arp2/3, 100 nM VCA, 1 μM Δmod, 1 μM V_1ab4_ and 0.6 μM VASP. The images show the reaction at 590 s. See supplementary movie 5. Scale bar = 15 μm. **(D)** Kymographs from (C) showing the elongation of a single actin filaments (top), actin filament branching in the presence of Arp2/3 + VCA (elongation rate ~9 su/s) (middle), and bundling in the presence of Arp2/3 + VCA supplemented with Δmod, V_1ab4_ and VASP (elongation rate ~15su/s) (bottom). (-) pointed end, (+) barbed end.

## Discussion

Our study shows how three proteins cooperate to adapt the architecture and dynamics of actin networks during the formation of adherens junctions (Figure 7A). Specifically, we showed that α-catenin, vinculin and VASP combine their activities to inhibit Arp2/3-mediated branching, stimulate the nucleation and elongation of linear actin filaments from profilin-actin and crosslink these filaments into bundles (Figure 7B, C).

**Figure 7.**
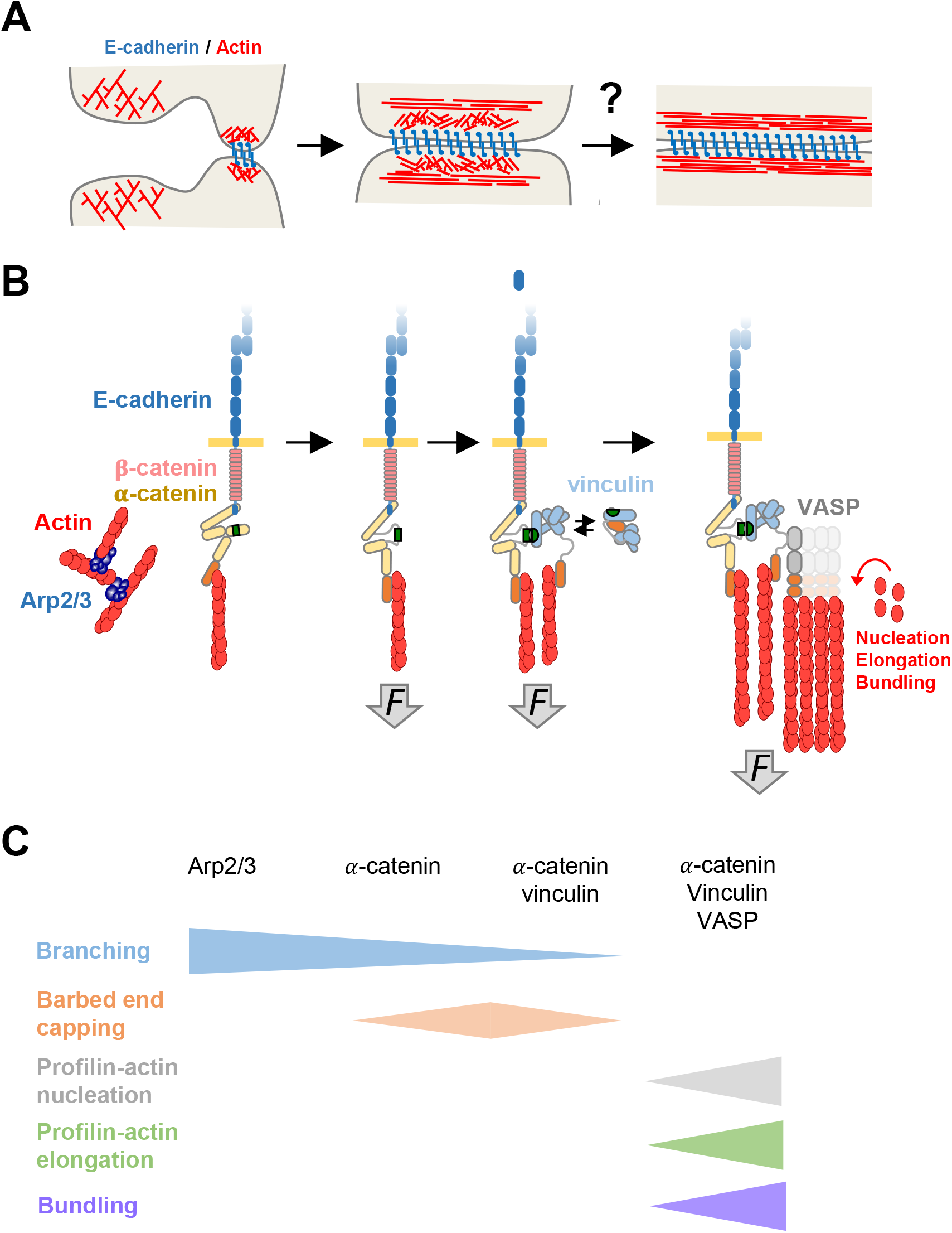
Mechanism by which α-catenin, vinculin and VASP regulate actin assembly during cellcell junction formation. **(A)** Transition from nascent to mature adherens junctions. **(B,C)** Regulation of actin assembly associated to cadherin-based cell-cell adhesions. **(C)** The color code correspond to the activities mentioned on the left side.

Isolated proteins have often been considered as functional entities and many studies have naturally sought to correlate their actin polymerization activities measured in vitro with actin dynamics observed in cells. In reality, many proteins are only parts of more complex machineries, as our study illustrates. Indeed, α-catenin was first described to cap actin filaments barbed ends (Hansen et al., 2013), while vinculin caps and nucleates filaments (Le Clainche et al., 2010), and VASP combines capping, nucleation and elongation activities (Breitsprecher et al., 2011, 2008; Hansen and Mullins, 2010; Laurent et al., 1999). Surprisingly we show here that the activity that results from their combination is not the sum of their individual activating and inhibiting activities. The mechanisms by which α-catenin, vinculin and VASP combine their activities to stimulate actin assembly seem to depend on the activities of VASP, such as nucleation and elongation which are enhanced by vinculin and α-catenin.

The mechanism of actin nucleation by VASP is not fully understood, but it can be hypothesised that the tetrameric state of this protein allows its monomeric actin binding domains (GABs) to bring actin monomers together to create an actin filament nucleus (Zimmermann et al., 2002; Chereau and Dominguez, 2006; Laurent et al., 1999; Ferron et al., 2007). It is possible that, in our in vitro assays, the formation of a complex, containing α-catenin dimers, vinculin dimers and VASP tetramers, results in complexes containing up to four VASP tetramers, further increasing the efficiency of actin nucleation. In cells, however, α-catenin forms heterodimers with β-catenin (Drees et al., 2005), preventing its homodimerization, and associates with a single vinculin dimer, which in turn associates with two VASP tetramers. This nucleation mechanism probably exists in cells because we observed it at different ionic strengths and in the presence of profilin.

The mechanism of actin filament barbed end elongation by VASP is well established as it requires the clustering of VASP tetramers, allowing the clusters of monomeric actin-binding domains (GABs) to feed actin monomers to the ends of filaments held by the actin-filament binding domains (FABs) (Breitsprecher et al., 2008, 2011; Brühmann et al., 2017). Bringing more VASP tetramers together through the formation of α-catenin-vinculin-VASP complexes is likely the reason for the increased elongation observed in our assays. In this process, VASP seems to protect the growing barbed ends from the capping activities of vinculin or alpha-catenin, as it protects against the action of the heterodimeric capping protein (CP) (Bear et al., 2002; Breitsprecher et al., 2008; Hansen and Mullins, 2010). However, our results cannot be explained by a simple competition between the three proteins for actin filament barbed ends since vinculin and α-catenin significantly increases the activity of VASP. Interestingly, the effect of vinculin and α-catenin, two bundling proteins, on VASP elongation activity is reminiscent of the effect of the bundling protein fascin on the elongation activity of the Drosophila Ena protein, a member of the Ena/VASP proteins (Winkelman et al., 2014), suggesting a general mechanism of stimulation of VASP elongation activity by bundling proteins.

Finally, our biochemical study must be placed in the context of cell-cell junctions. In cells, the mechanism of junction formation involves actin polymerization, which has been proposed to promote lateral clustering of cadherins (Truong Quang et al., 2013), while the tensile force of actomyosin would strengthen the junctions (Yonemura et al., 2010). More recent studies in kidney epithelial cell sheets (MDCK cells) indicate that cadherin puncta observed in cells in fluorescence microscopy correspond to interdigitations of the plasma membrane (Li et al., 2021). These interdigitations are membrane protrusions whose formation requires actin polymerization under the control of the Arp2/3 complex, EVL and CRMP-1 (Li et al., 2020). Electron microscopy experiments revealed that two distinct actin network are associated to AJs; a juxtamembrane branched actin network and a perijunctional bundle of filaments parallel to the membrane (Efimova and Svitkina, 2018) (Figure 7A). Interestingly, junctional myosin isoforms have specific localization and functions, as NMIIA is associated with actin bundles to generate force, while NMIIB is associated with the branched network to promote force transmission (Heuzé et al., 2019). Other studies have shown that the α-catenin-vinculin-VASP complex induces the polymerization of actin filaments at AJs in response to the force generated by actomyosin (Leerberg et al., 2014). Our results suggest that, in response to actomyosin force, the branched actin network associated with AJs is also reorganized into bundles. Consistent with this hypothesis, NMIIB knock-down cells exhibit AJs that contain less open α-catenin and are associated with thicker Arp2/3-containing actin networks (Heuzé et al., 2019). However, to date there have been no comparative electron microscopy studies of the ultrastructure of actin networks associated with AJs subjected to increasing forces to test this hypothesis.

In addition to Arp2/3 and α-catenin-vinculin-VASP machineries, formins, such as Dia-1 and formin-1, known to promote the elongation of unbranched actin filaments (Romero et al., 2004), play a role in actin assembly at AJs (Acharya et al., 2017; Carramusa et al., 2007; Kobielak et al., 2004). Further studies are needed to understand the organization and dynamics of actin networks that result from coordination between the activities of Arp2/3, formins, α-catenin-vinculin-VASP, and the myosins NMIIA and NMIIB.

## Materials and methods

### cDNA constructs

α-catenin WT and Δmod cDNA were cloned into a pDW363 plasmid with a C-terminal His6 tag (Seddiki & al., 2017). cDNA encoding for vinculin E28K/D33H/D110H/R113E/N773I/E775K (V_1ab4_) was synthesized and subcloned in the NcoI site of pET-3d by Genscript, with a C-terminal His8 tag (Wang et al., 2022). The construct was verified by sequencing. The cDNA encoding for human full-length VASP was cloned into a pGEX-6P1 plasmid to produce a N-terminal cleavable Glutathione-S-transferase (GST)-fused VASP.

### Protein purification

Rabbit skeletal muscle actin was purified from acetone powder (Wiesner, 2006). Cycles of polymerization and depolymerization were followed by dialysis in 5 mM Tris pH 7.8, 0.2 mM ATP, 0.1 mM CaCl2, 0.01% NaN3, 1 mM DTT, and gel filtration on a Superdex G-200 column (GE Healthcare). Actin was labelled with N-pyrenyliodoacetamide, and Alexa Fluor 488 (Invitrogen) or Atto 488 (Atto-TEC) Succinimidyl Ester as previously described (Ciobanasu et al., 2015). Arp2/3 and profilin were purified as previously described (Le Clainche and Carlier, 2004). Spectrin-actin seeds were prepared as previously described (Casella et al., 1986).

Expression of all the recombinant proteins in this study was performed with the same protocol. After plasmid transformation in *Escherichia coli* (BL21 DE3, Invitrogen), bacteria were grown in 3-6 liter of LB medium containing 0.1 mg.ml^-1^ of appropriate antibiotics (ampicillin or kanamycin) at 37°C until the absorbance reached 0.6-0.8 at 600 nm. 1 mM of isopropyl 1-thio-β-D-galactopyranoside (IPTG) was added to the medium to induce the expression of the recombinant proteins of interest during 16h at 16°C. After centrifugation, the α-catenin WT, α-catenin Δmod and VASP FL bacterial pellets were lysed by sonication in 20 mM Tris pH 7.8, 500 mM NaCl, 1 mM β-mercaptoethanol, 10 μg.ml^-1^ benzamidine and 1 mM PMSF. The V_1ab4_ bacterial pellets were lysed by sonication in 20 mM Tris pH 8.0, 1 M NaCl, 1 mM β-mercaptoethanol, 10 μg.ml^-1^ benzamidine and 1 mM PMSF.

Lysates of His-tagged α-catenin WT and Δmod were bound to Ni-NTA (Ni^2+^-nitrilotriacetic acid)-sepharose affinity chromatography (Qiagen), washed with 50 mM Tris pH 7.8, 500 mM NaCl, 20 mM imidazole, 1 mM β-mercaptoethanol, eluted with 50 mM Tris pH 7.8, 500 mM NaCl, 250 mM imidazole, 1 mM β-mercaptoethanol, and purified on a gel filtration column (Superdex 200, 16/60, GE Healthcare). α-catenin constructs were finally dialyzed in 20 mM Tris pH 7.8, 150 mM KCl, 1 mM DTT, frozen in liquid nitrogen and stored at - 80°C.

V_1ab4_ was purified by Ni-NTA-sepharose affinity chromatography (Qiagen), followed by a Q-Sepharose ion exchange column. V_1ab4_ protein was finally dialyzed in 20 mM Tris pH 7.8, 1 mM DTT, frozen in liquid nitrogen and stored at −80°C.

Human full-length VASP lysates were first bound to Glutathione Sepharose in 50 mM Tris pH 7.8, 500 mM NaCl, then cleaved from GST by Prescission protease. The protein was dialyzed in 20 mM Tris, 100 mM KCl, 1 mM DTT, frozen in liquid nitrogen and stored at - 80°C.

### Actin polymerization assay

Actin polymerization was measured by the increasing in fluorescence of 10% pyrene-labelled actin. For barbed-end elongation measurements, 100 pM of spectrin-actin seeds were added to the reaction and actin polymerization was induced by adding 100 mM KCl, 1 mM MgCl2 and 0.2 mM EGTA to a solution of 10% pyrene-labeled Ca-ATP-G-actin containing the proteins of interest. To test the nucleation activity of proteins, spontaneous polymerization was induced by adding 25 mM KCl, 1 mM MgCl2 and 0.2 mM EGTA to a 10% pyrene-labeled Ca-ATP-G-actin solution in the presence of proteins of interest. Fluorescence measurements were carried out with a Safas Xenius FLX spectrophotometer (Safas, Monaco). The graphs and plots were assembled with Excel or Kaleidagraph. The experiments were reproduced 2 to 3 times with the same conclusions.

### Light scattering measurements and microscopy observation of actin bundling

The actin filaments bundling during actin polymerization was examined by measuring the light scattering at 400 nm with a Safas Xenius FLX spectrophotometer (Safas, Monaco). Actin polymerization was induced by adding 25 mM KCl, 1 mM MgCl2 and 0.2 mM EGTA to a solution of 2% Alexa488-labeled-G-actin in the presence of the proteins of interest. A 5 μl sample was taken from the reaction after 1500 s for epifluorescence microscopy observation. The graphs and plots for light scattering kinetics were assembled with Excel or Kaleidagraph. For epifluorescence microscopy, the images were acquired with an Olympus IX71 inverted microscope equipped with a 60X oil immersion objective (Olympus, 1.45 NA) and coupled to an EMCCD camera (Cascade, Photometrics). Images were acquired with MicroManager and then assembled with ImageJ software. The experiments were reproduced 3 times with the same conclusions.

### Observation of single actin filaments in TIRF microscopy

Our protocol is based on a modification of previously published methods (Ciobanasu et al., 2018; Le Clainche et al., 2010). After sonication with milliQ water and ethanol, the coverslips (24 mm x 40 mm, Thermo Scientific/Menzel-Glaser) were irradiated for 1 min under a deep UVs lamp (Ossila), and then incubated with 0.1 mg.ml^-1^ PLL-g-PEG (SuSoS) dissolved in 10 mM HEPES pH 7.4 for 1 h at room temperature. The coverslips were then washed with milliQ water and dried. Flow chamber, with a typical volume of 50 - 70 μl, was created by sticking the PLL-PEG passivated side of the coverslip to a glass slide (Super Frost, Thermo Scientific) using a double-sided adhesive tape. After incubation with washing buffer (5 mM Tris pH 7.8, 200 μM ATP, 1 mM DTT, 1 mM MgCl2, 25 or 100 mM KCl) for 1 min, the chamber was saturated with 10% bovine serum albumin (BSA) for 5 min, and then washed with washing buffer. A final reaction composed of 1.5 μM G-actin (10% Alexa Fluor or Atto 488-labeled) in 5 mM Tris pH 7.8, 200 μM ATP, 0.4% methylcellulose, 5 mM 1,4-diazabicyclo(2,2,2)-octane (DABCO), 100 mM KCl or 25 mM KCl, 1 mM MgCl2, 200 μM EGTA, 10 mM DTT supplemented with our proteins of interest, was injected in the chamber. The chamber was sealed with oil or VALAP (mix of vaseline, lanolin and paraffin), and then observed on an Olympus IX71 inverted microscope equipped with a 60X oil immersion objective (Olympus, 1.45 NA) and coupled to an EMCCD camera (Cascade, Photometrics). The time-lapse videos were acquired with MicroManager and analyzed by ImageJ. The plots were assembled with Excel or Kaleidagraph. All the experiments were reproduced 3 times with the same conclusions. Statistical analyses were carried out with a Mann-Whitney non parametric test in Microsoft Excel.

## Acknowledgements

This work was supported by the Agence Nationale de la Recherche grants ANR-18-CE13-0026-01 RECAMECA to CLC, ANR-20-CE13-0016 MECOLLACT to CLC and AMG, ANR-21-CE13-0010-03 MYOCORTEX to CLC. HW is supported by a PhD fellowship from the China Scholarship Council (CSC). RS is supported by a PhD fellowship from Association de Spécialisation et d’Orientation Scientifique. The present work has benefited from the Light Microscopy facility of Imagerie-Gif, (http://www.i2bc.paris-saclay.fr), member of IBiSA (http://www.ibisa.net), supported by “France-BioImaging” (ANR-10-INBS-04-01), and the Labex “Saclay Plant Sciences” (ANR-10-LABX-0040-SPS). We thank the members of the “Cytoskeleton Dynamics and Motility” team for helpful discussions.

---

**Supplementary Figure 1.**
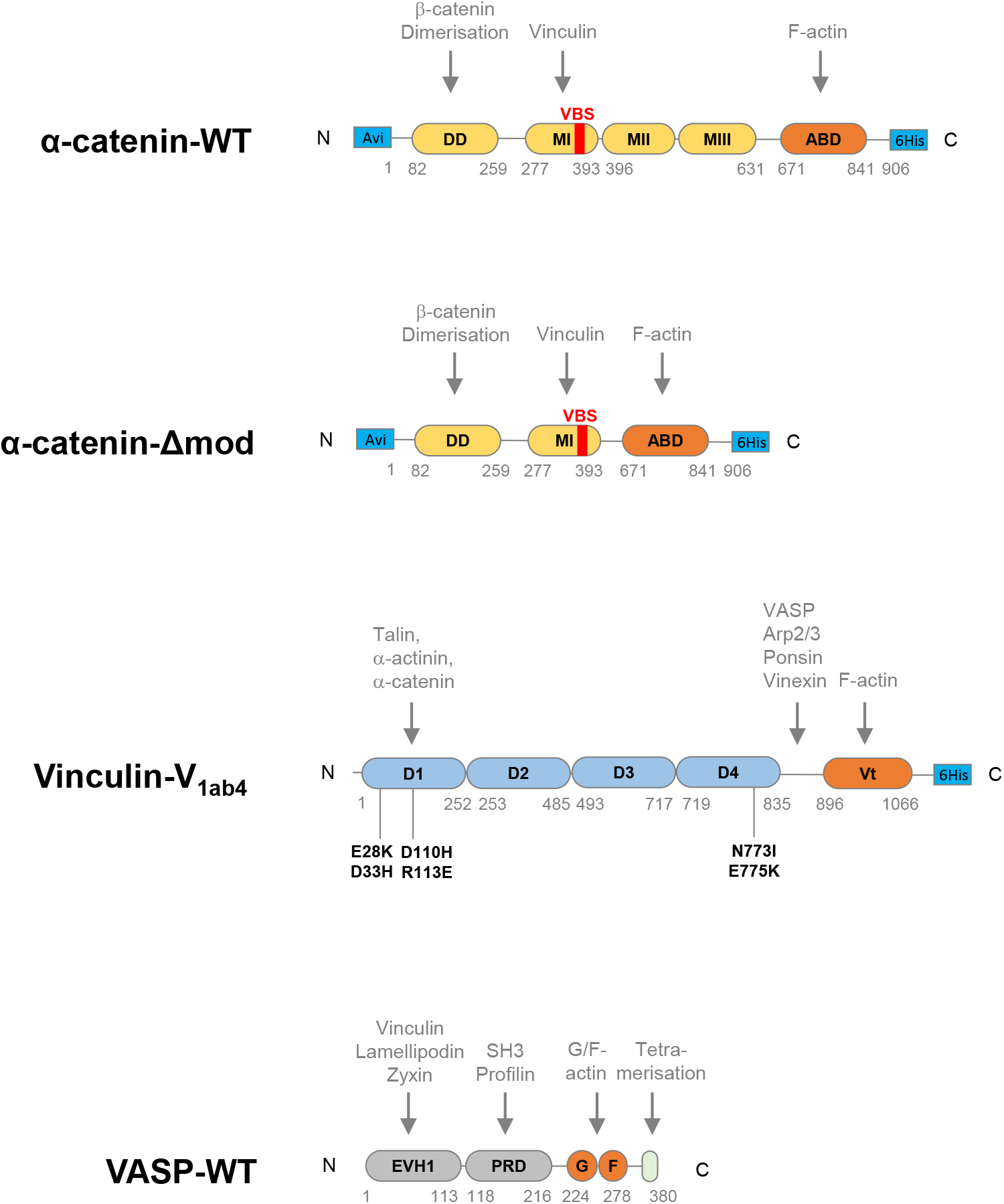
Schematic representation of proteins constructs used in this study. DD, dimerization domain; MI-MII-MIII, modulator domain; ABD, actin binding domain (orange); VBS, vinculin binding site (red); Vt, vinculin tail; EVH1, Ena/VASP homology domain 1; PRD, proline-rich domain; E28K/D33H, D110H/R113E, N773I/E775K, point mutations in vinculin-V_1ab4_ mutant. Proteins partners shown in gray can link to the corresponding domain of each protein of interest.

**Supplementary Figure 2.**
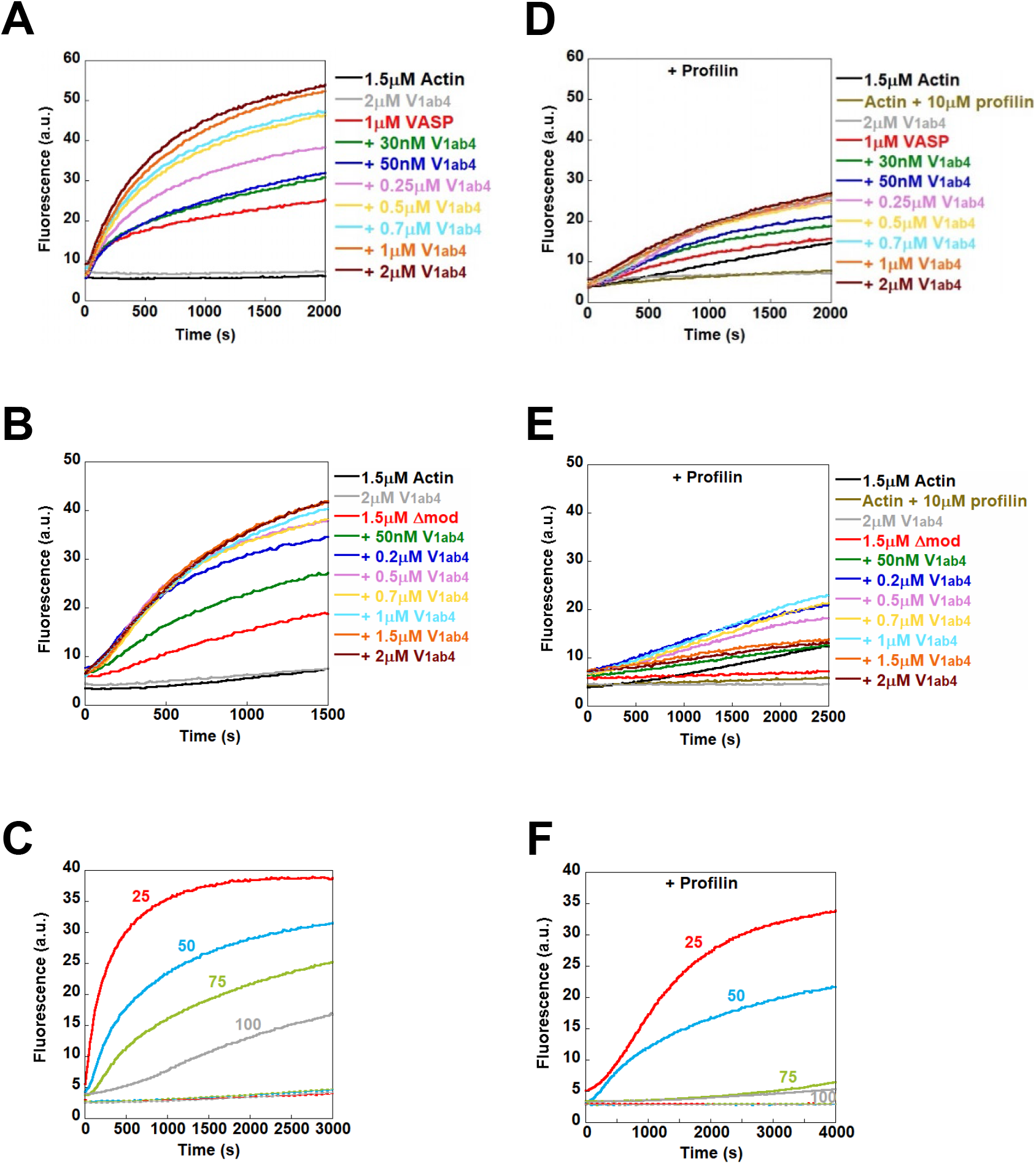
Vinculin V_1ab4_ synergizes with α-catenin Δmod or VASP to stimulate actin polymerization. **(A,B)** Spontaneous actin polymerization (1.5 μM G-actin, 10% pyrene-labeled) was measured in the presence of increasing concentrations of V_1ab4_ in the presence of 1 μM VASP **(A)** or 1.5 μM Δmod **(B)**. **(C)** Spontaneous actin polymerization (1.5 μM G-actin, 10% pyrene-labeled) was measured in the presence of increasing concentrations of KCl in the presence of 1.5 μM Δmod, 2 μM V_1ab4_ and 1 μM VASP. **(D,E)** Spontaneous actin polymerization (1.5*μ*M G-actin, 10% pyrene-labeled) was measured in the presence of 10 μM profilin and increasing concentrations of V_1ab4_ in the presence of 1*μ*M VASP **(D)** or 1.5 μM Δmod **(E)**. **(F)** Spontaneous actin polymerization (1.5 μM G-actin, 10% pyrene-labeled) was measured in the presence of increasing concentrations of KCl, 10 μM profilin,1.5 μM Δmod, 2 μM V_1ab4_ and 1 μM VASP. The dashed lines in (C) and (F) correspond to the control with actin alone at 25, 50, 70 and 100 mM KCl.

**Supplementary Figure 3.**
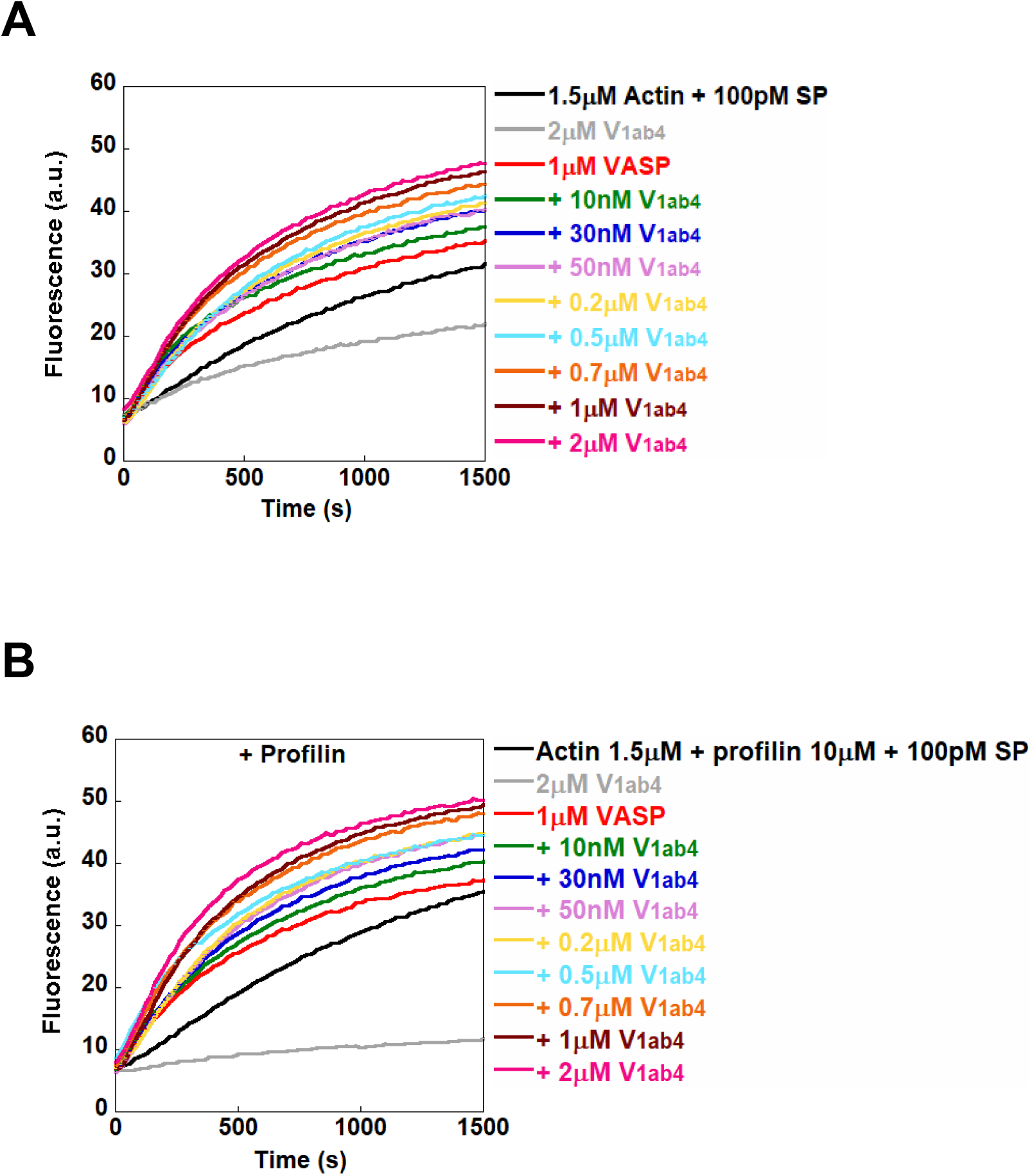
Elongation of actin filament barbed ends by VASP in the presence of V_1ab4_. **(A,B)** The elongation of actin filament barbed ends was measured in the presence of 100 pM spectrin-actin seeds (SP), 1.5 μM G-actin (10% pyrene-labeled), 1 μM VASP, and increasing concentration of V_1ab4_ in the absence **(A)** or the presence of 10 μM profilin **(B)**.

## Legends of supplementary movies

**Supplementary Movie 1. Observation of the α-catenin-vinculin-VASP activity on single actin filaments in TIRF microscopy at 25 mM KCl.**

Conditions: 0.8 μM G-actin (10% atto 488-labeled) in fluorescence buffer (5 mM Tris pH7.8, 200 μM ATP, 5 mM 1,4-diazabicyclo(2,2,2)-octane (DABCO), 0.4% methylcellulose, 25 mM KCl, 1 mM MgCl_2_, 200 μM EGTA, 20 mM DTT), supplemented with 1 μM Δmod or 1 μM V_1ab4_ or 0.6 μM VASP or the combination of the three proteins together. Scale bar = 15 μm.

**Supplementary Movie 2. Observation of the α-catenin-vinculin-VASP activity on single actin filaments in TIRF microscopy at 100 mM KCl.**

Conditions: 0.8 μM G-actin (10% atto 488-labeled) in fluorescence buffer (5 mM Tris pH7.8, 200 μM ATP, 5 mM 1,4-diazabicyclo(2,2,2)-octane (DABCO), 0.4% methylcellulose, 100 mM KCl, 1 mM MgCl_2_, 200 μM EGTA, 20 mM DTT), supplemented with 1 μM Δmod or 1 μM V_1ab4_ or 0.6 μM VASP or the combination of the three proteins together. Scale bar = 15 μm.

**Supplementary Movie 3. Observation of the α-catenin-vinculin-VASP activity on single actin filaments in TIRF microscopy in the presence of profilin, at 25 mM KCl.**

Conditions: 0.8 μM G-actin (10% atto 488-labeled) in fluorescence buffer (5 mM Tris pH7.8, 200 μM ATP, 5 mM 1,4-diazabicyclo(2,2,2)-octane (DABCO), 0.4% methylcellulose, 25 mM KCl, 1 mM MgCl_2_, 200 μM EGTA, 20 mM DTT), supplemented with 5 μM profilin alone and in the presence of 1 μMΔmod, 1 μM V_1ab4_ and 0.6*μ*M VASP. Scale bar = 15 μm.

**Supplementary Movie 4. Observation of the α-catenin-vinculin-VASP activity on single actin filaments in TIRF microscopy in the presence of profilin, at 100 mM KCl.**

Conditions: 0.8 μM G-actin (10% atto 488-labeled) in fluorescence buffer (5 mM Tris pH7.8, 200 μM ATP, 5 mM 1,4-diazabicyclo(2,2,2)-octane (DABCO), 0.4% methylcellulose, 100 mM KCl, 1 mM MgCl_2_, 200 μM EGTA, 20 mM DTT), supplemented with 5 μM profilin alone and in the presence of 1 μM Δmod, 1 μM V_1ab4_ and 0.6 μM VASP. Scale bar = 15 μm.

**Supplementary Movie 5. Remodeling of Arp2/3-mediated branched actin into fast-growing bundles by α-catenin, vinculin and VASP.**

Conditions: 0.8 μM G-actin (10% atto 488-labeled) in fluorescence buffer (5 mM Tris pH7.8, 200 μM ATP, 5 mM 1,4-diazabicyclo(2,2,2)-octane (DABCO), 0.4% methylcellulose, 50 mM KCl, 1 mM MgCl_2_, 200 μM EGTA, 20 mM DTT), supplemented with the indicated combinations of 50 nM Arp2/3, 100 nM VCA, 1 μM Δmod, 1 μM V_1ab4_ and 0.6 μM VASP. Scale bar = 15 μm.

